# The effect of spatial variation for predicting aphid outbreaks

**DOI:** 10.1101/367953

**Authors:** Christian Damgaard, Marianne Bruus, Jørgen Aagaard Axelsen

## Abstract

In order to improve forecasting of pest epidemics, it is important to know the spatial scale at which specific forecasts are reliable. To investigate the spatial scale of aphid outbreaks, we have developed a spatio-temporal stochastic aphid population growth model, and fitted the model to empirical spatial time-series aphid population data using a Bayesian hierarchical fitting procedure. Furthermore, detailed spatial data of the initial phases of population growth was investigated in a semivariogram. Our results suggest that there is limited spatial variation in the initial occurrence probability at a spatial scale of 10 km. Consequently, the results support the hypothesis that initial aphid population sizes and outbreaks may be predicted in fields within a 10 km radius. For farmers, this may imply that they can rely their decision of whether to spray against aphids on observations made by other nearby farmers or by the consultancy service.

## Introduction

Insect pests are responsible for serious negative effects on food production. The common cereal aphids, the Cherry oat aphid (*Rhopalosiphum padi*), the Grain aphid (*Sitobion avenae*) and the Rose-grain aphid (*Metopolophium dirrhodum*), are known to cause considerable losses to winter wheat in large parts of the world, and these species have been estimated to cause losses of 700.000 tons per year in Europe due to direct damage (Wellings et al. 1989). On top of this figure comes the indirect damage due to virus transmission and sooty mold caused by the excretion of honeydew (Larsson 2005). Therefore, there is a great interest in controlling aphids in winter wheat, and a number of simulation models have been developed to predict the population development of these pests (e.g. Ciss et al. 2014; Duffy et al. 2017; Holst & Ruggle 1997; Klueken et al. 2009). Furthermore, some population dynamic models have been combined with modules on economy and pesticide choices to produce decision support systems with the aim of helping farmers optimizing the timing of pesticide applications. Such decision support systems include CPO in Denmark (Hagelskjær & Jørgensen 2003) and GETLAUS (Gosselke et al. 2001) in Germany. In a review of decision support systems, (Axelsen et al. 2012) concluded that farmers generally do not use decision support systems, and one of the reasons is that they do not find it worth the effort to spend time on estimating aphid input densities. Instead, many farmers perform precautionary insecticide treatments, which may not be economically sound.

The grain aphid is the most numerous aphid in winter wheat (Hansen 2003) in Denmark. In temperate regions, it overwinters on grasses as eggs. When temperatures rise in spring, the eggs hatch into unwinged females, which reproduce parthenogenetically. In the second or third generation, the majority of aphids are winged individuals, which migrate into the cereal fields. The formation of winged individuals is initiated by a complex combination of crowding (i.e. high densities), light and temperature (Hansen 2003). Despite high temperatures, crowding may be prevented by adverse temperatures, precipitation, predators and parasites. Once in the cereal fields, temperature is the key driver for aphid population growth, as shown by Jones (1979). However, population growth may be modified by heavy rainfall, wind, parasites, disease and predators (Hansen 2003; Jones 1979; Nakata 1995). Plantegenest et al. (2001) showed that not including the effects of parasotoids and especially fungal diseases may cause over-estimates of aphid population growth. Wang et al. (2015) and Chabert & Sarthou (2017) found that fertilizer input may increase aphid populations. A few papers have studied the impact of heavy rainfall on aphid populations. In cotton fields, Chamuene et al. (2018) found that in some, but not all cases rain incidents of less than 50 mm significantly increased the mortality of the aphid *Aphis gossypii*. A nine-year study of soybean fields found significant effects of precipitation patterns on the population dynamics of *Aphis glycines* (Stack Whitney et al. 2016). In laboratory experiments, Mann et al. (1995) found that the loss of *Sitobion avenae* from oat plants was correlated with rain intensity and duration, and increased when combined with wind gusts. Jones (1979) observed that heavy rainfall before the ears appear may reduce aphid numbers on cereals, and that information about rainfall may complement the prediction of population size based on temperature.

In order to optimize the decision support systems, large efforts have been spent on improving model performance in relation to weather parameters and natural regulation. These efforts have without doubt improved the models, but taking into consideration that aphid populations develop exponentially, and that the population can double in 55 hours at 20°C (Dedryver et al. 2010), it is critical that the initial densities are determined at high precision in order to predict the epidemic development. However, surprisingly little efforts have been devoted to obtaining quick and reliable estimates of the initial densities, although several methods have been developed to estimate the density of aphids in cereals. For instance, Elliott et al. (1990) developed binomial sequential sampling plans that require rather larger efforts at low densities to produce reliable estimates, and Hansen (1991), who suggested to investigate aphid presence/absence on 50 or 100 tillers and used an equation to convert to aphids per tiller. Both methods take some time, and come up with assessments of average densities with some uncertainties. The uncertainties are predefined in the sequential sampling plan and the required uncertainty level is decisive for the number of plants to investigate. When counting presence/absence on a number of straws, the uncertainty can be calculated before being used in simulation models and decision support systems. However, none of the models and decision support systems appear to use uncertainty in their projections of aphid population development, and in turn relate uncertainties to the output. Given that aphids show exponential population growth, uncertainties in the estimate of initial densities can cause large uncertainties to the projections of the aphid population density some weeks later. This uncertainty should ideally be reflected in the suggestions produced by a decision support system.

To predict the spatial and temporal development of aphid outbreaks it is important to sample and model population data that encompasses both spatial and temporal dimensions. In this study, we have sampled aphid populations in a spatial setup during both the initial and the epidemic phase to fit a population growth model. More specifically, we have developed a spatio-temporal stochastic aphid population growth model and fitted the model to empirical spatial time-series aphid population data using a Bayesian hierarchical fitting procedure. Such Bayesian hierarchical population models have successfully been applied on a number of pest cases, e.g. the population dynamics of coffee berry borer infestation (Ruiz-Cárdenas et al. 2009). The fitted spatio-temporal population growth model may be used to generalize existing deterministic aphid forecasting models with the effect of stochastic spatial variation. Furthermore, we use the fitted model and complementary spatial statistics to investigate the hypothesis that initial aphid population sizes and outbreaks may be predicted in fields within a 10 km radius of the nearest aphid-monitoring site.

## Materials and Methods

### Field sites and aphid sampling

The occurrence probability of the grain aphid, *Sitobion avenae*, i.e. the percentage wheat straws with aphids (correlated with both number of aphids per straw, and number of aphids per area (Feng et al. 1993; Hansen 2003; Hein et al. 1995)), was recorded in twelve wheat fields in central Jutland, Denmark, in 2016 and 2017. The wheat fields (=sites) were laid out in hierarchical geographic design with three regions of four sites. In Denmark, aphids are monitored in the official aphid-sampling programme (Observation Web, https://www.landbrugsinfo.dk/planteavl/plantevaern/varslingregistrerings-net/-sider/startside.aspx). In the current project, we used an Observation Web field as centre site for each of the three regions, and within each region, three other sites (fields), positioned approximately three, six and ten kilometers away (Fig. 1), were studied. As a consequence of the proximity of some of the Observation Web fields, the three regions overlap to some extent. At each site, the occurrence probability of aphids on individual plants were recorded and the occurrence probability was used as a proxy for the aphid population size, based on the fact that percentage straws with aphids is highly correlated with the number of aphids per square meter (Feng et al. 1993; Hansen 2003; Hein et al. 1995).

**Fig. 1.**
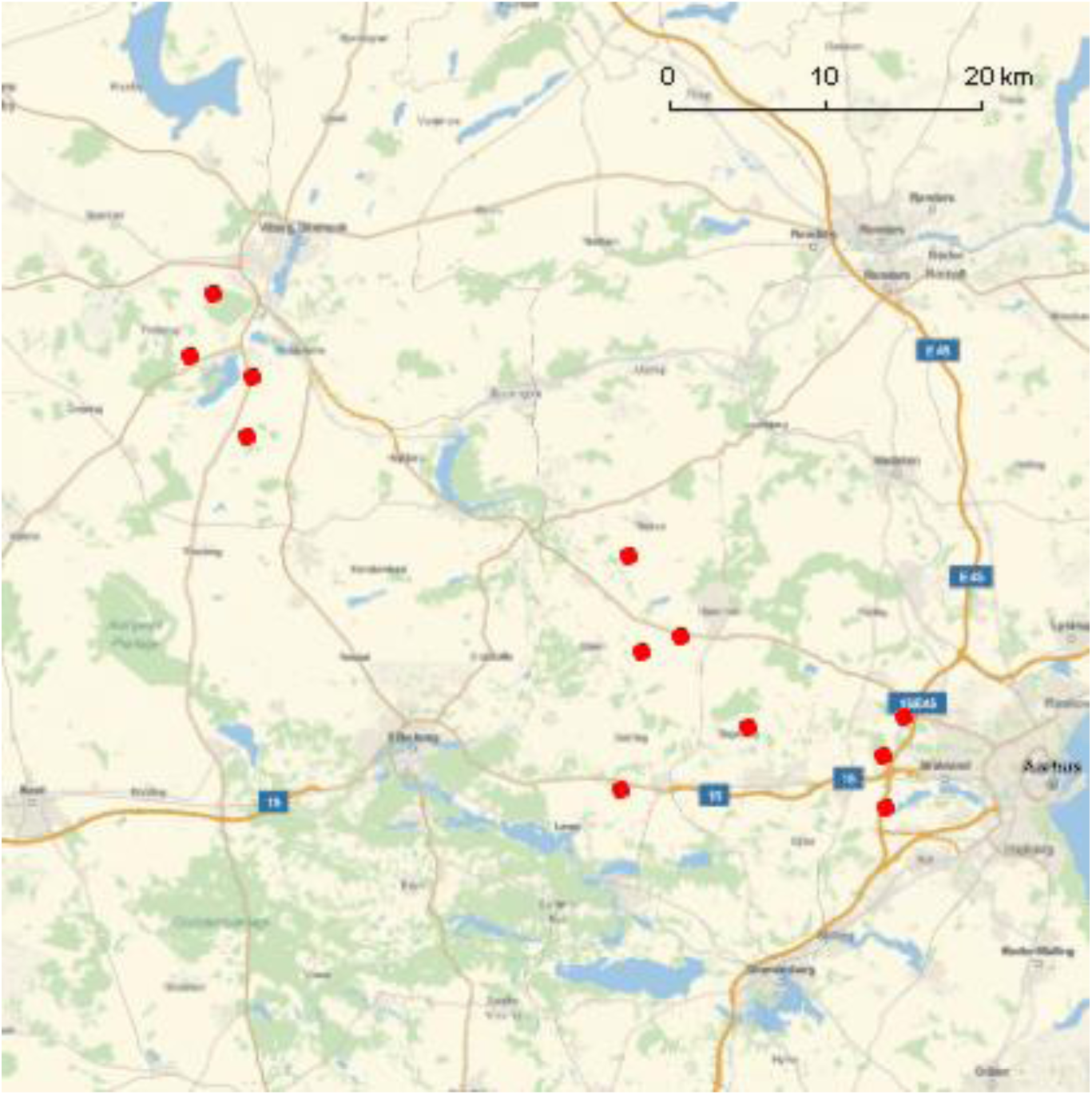
Twelve wheat fields in the middle of Jutland, Denmark, where aphids were sampled. The wheat field were laid out in hierarchical geographic design with three regions of four fields. Each region had a center field with fields positioned approximately three, six and ten kilometers away.

In 2016, aphid occurrence was recorded for five samples of either 80 or 100 wheat plants at each site on May 24, June 6, June 13, June 20, June 27, July 4, and July 11. The five samples within a site were taken along a transect with at least 50m between samples. Site-specific degree-days were calculated from the average day temperature at the weather station closest to each site with a base temperature of 5 degrees Celsius. Furthermore, the intensity of precipitation events at the different sites was recorded daily by a weather station placed at each field (Agrimex® Rosenborg 35980).

In order to complement the result of the spatial variation of the initial phases of aphid outbreaks obtained in the spatial modelling of the aphid occurrence data sampled in 2016, aphid occurrence was recorded more intensively at the same twelve sites in the beginning of the growth period the following year and analyzed in a semivariogram. In 2017, aphid occurrence was recorded for ten samples of 50 wheat plants at each site on May 30 and June 7. The ten samples within a site were laid out along three transects with at least 50m between all plots, and the exact geographical position of each sample was determined.

### Statistical modelling of the aphid population

The spatio-temporal aphid occurrence data is modelled using Bayesian hierarchical methods (Clark & Gelfand 2006). The observed number of straws with at least one aphid at site *i* and plot *k* at degree-day *t* is denoted *y*_*i,k,t*_ and is assumed to be binomially distributed with *n*_*i,k,t*_, the number of straws sampled, and *p*_*i,t*_, the occurrence probability that a straw has at least one aphid at site *i* at degree-day *t*,

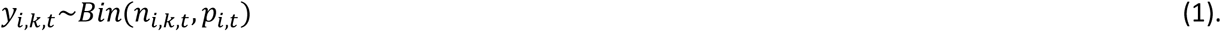

The site-specific occurrence probability is modelled using an exponential function of degree-day *t*,

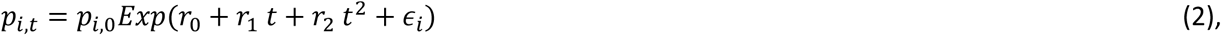

where *p*_*i,*0_ is the occurrence probability on a fixed initial day, *r*_0_, *r*_1_, and *r*_2_ are population growth parameters, and *ϵ*_*i*_ are Gaussian distributed site-specific random effects, 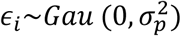.

The *n* site-specific initial occurrence probabilities are assumed to arise from a Gaussian process model,

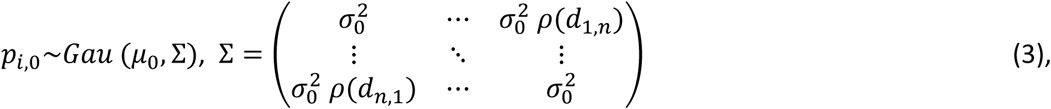

where *μ*_0_ is the mean initial occurrence probability, 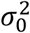 is the variance, and 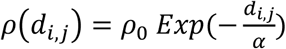 with *d*_*i,j*_ being the distance between site *i* and site *j*, *α* is the scale of the spatial effect that is set to 10 km, and *ρ*_0_ is a parameter that measures the spatial covariance (Haran 2011; Ovaskainen et al. 2016). The covariance matrix by definition has to be positive definite, which puts upper and lower bounds on *ρ*_0_.

The prior distributions of all parameters were assumed to be uniform except the two scale parameters that were assumed to be inverse gamma distributed, *σ*_*p*_∼*Invgam*(0.1,0.1) and *σ*_0_∼*Invgam*(0.1,0.1). The joint Bayesian posterior distribution of the parameters and the two latent variable vectors, *p*_*i,*0_ and *ϵ*_*i*_, each of dimension *n*, were calculated using Markov Chain Monte Carlo (MCMC), Metropolis-Hastings, simulations with a multivariate normal candidate distribution and using a MCMC run of 300,000 iterations after a burn-in period.

Plots of the sampling chains of all parameters and latent variables were inspected in order to check the mixing properties of the used sampling procedure, and whether the burn-in period was sufficient. Additionally, the overall fitting properties of the model were checked by inspecting the regularity and shape of the marginal distribution of all parameters as well as the distribution of the deviance (= –2 log*L*(*Y*|*θ*)). The efficiency of the MCMC procedure was assessed by inspecting the evolution in the deviance.

Statistical inferences on the parameters were based on the marginal posterior distribution of the parameters.

All calculations were done using *Mathematica* version 10 (Wolfram 2015).

## Results

The observed spatio-temporal mean occurrence probability of *S. avenae* in 2016 (Fig. 2) was fitted to the spatial growth model. The burn-in period of the MCMC was relatively long (600,000 iterations), but after the deviance had stabilized, the fitting properties of the models were judged to be acceptable based on visual inspections of the mixing properties of the parameters and the latent variables.

**Fig. 2.**
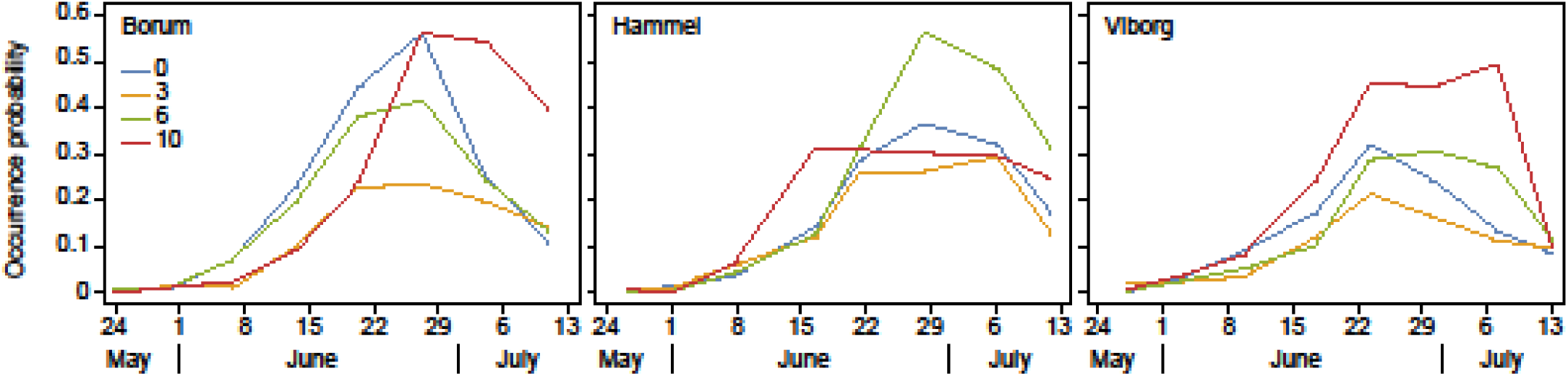
Observed spatio-temporal mean occurrence probability of *S. avenae.* The sites are located in three different regions (Borum, Hammel and Viborg). Each region had a center field (0) with fields positioned approximately three (3), six (6) and ten (10) kilometers away.

The deterministic part of the population growth model (eqn 1.) seemed to adequately model the dynamics of aphid occurrence probabilities as a function of degree days from June 1 2016 to July 1 2016 (Fig. 3) when shape and mode of the expected outbreak was visually compared to the observed spatio-temporal mean occurrence probability data in the same period (Fig. 2). Furthermore, there were no apparent pattern among the latent variables that model the among-site variation in population growth, *ϵ*_*i*_ (Fig. 4), which indicate that where were no systematic regional effects influencing growth apart from degree days.

**Fig. 3.**
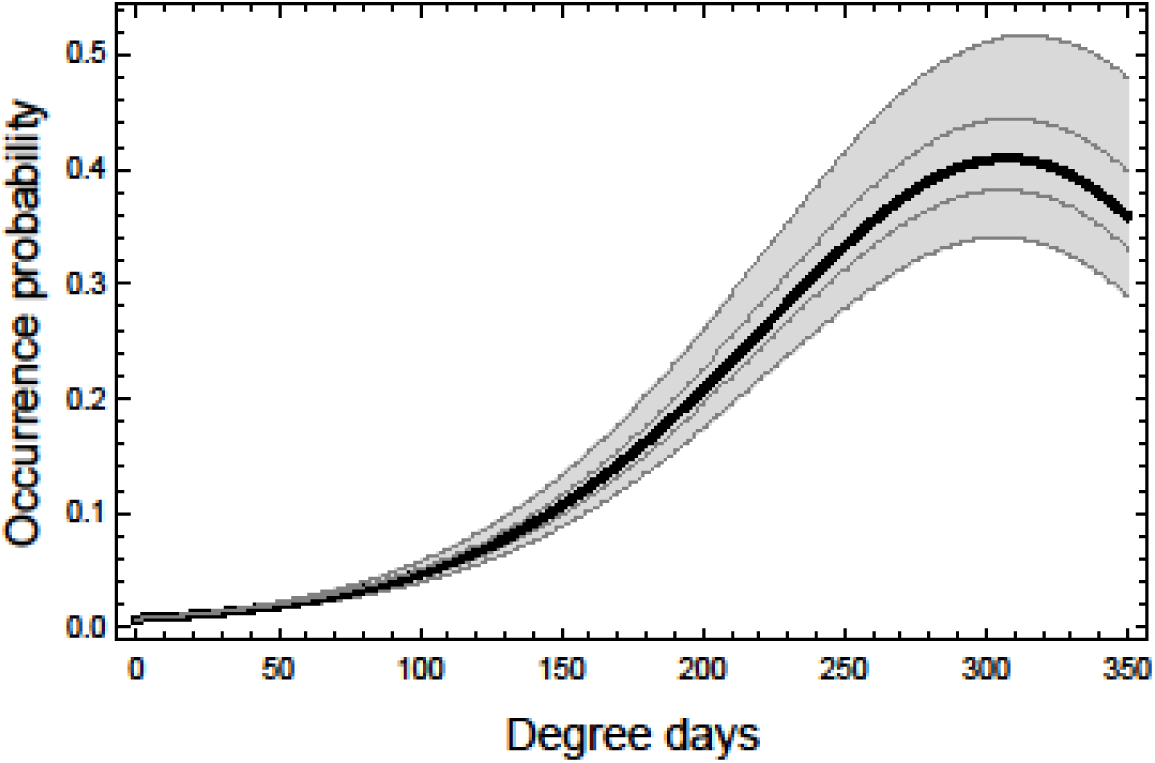
Expected occurrence probability of *S. avenae* at the mean degree-days from June 1 2016 to July 1 2016. The curve is calculated using equation (2), the expected *μ*_0_, and the joint posterior distribution of *r*_0_,*r*_1_, and *r*_2_. The thick line is the median occurrence probability and the two thin lines are the 25% percentile and 75% percentile of the occurrence probability. The shaded area is the 95% credibility interval of the occurrence probability.

**Fig. 4.**
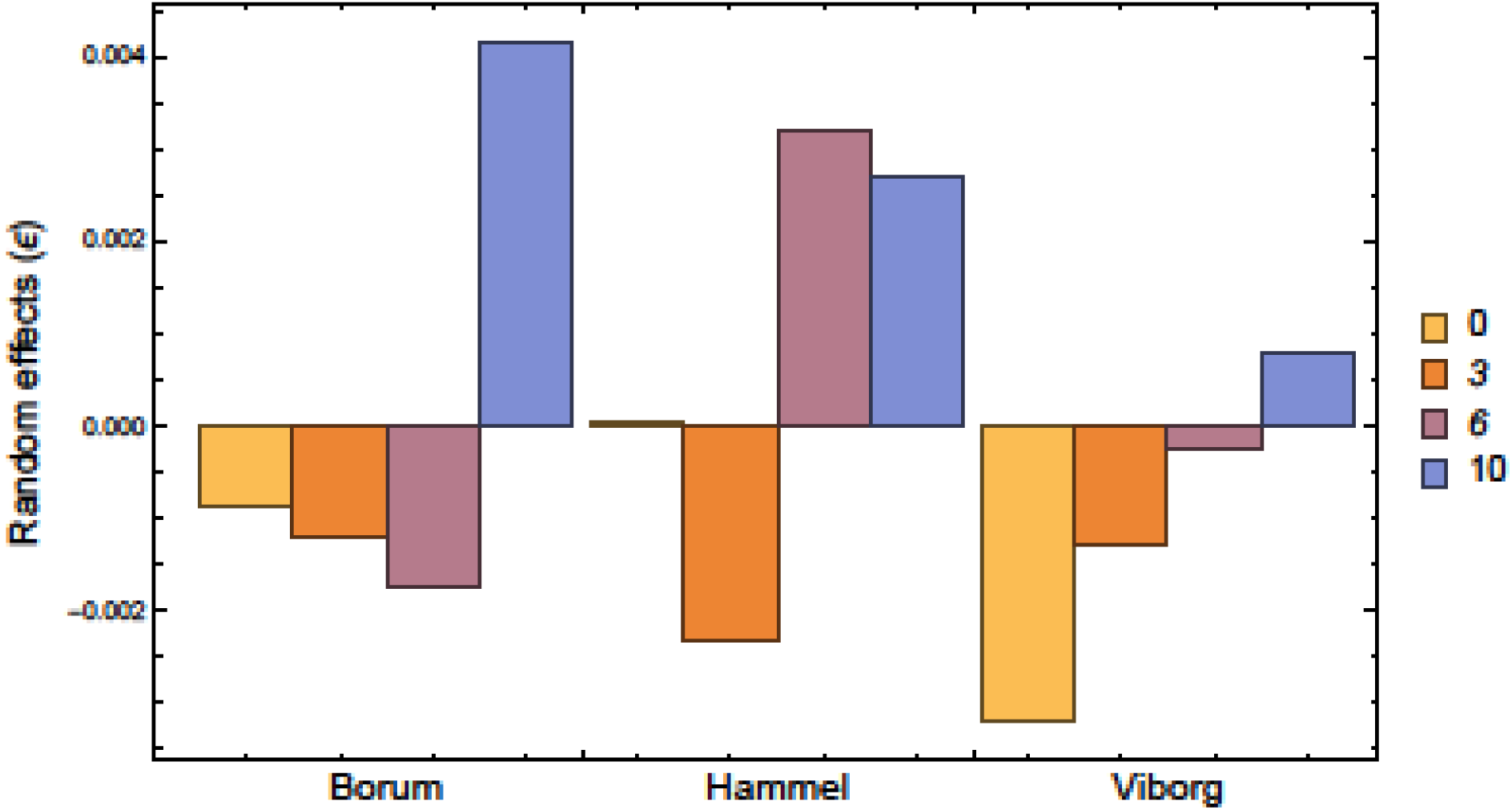
The distribution of the latent variables that model the among-site variation in population growth, *ϵ*_*i*_, at the different sites.

The marginal posterior distributions of the parameters of interest are summarized in Table 1 by their percentiles. The parameters were generally uncorrelated, except for, as expected, the estimates of the population growth parameters *r*_0_, *r*_1_ and *r*_2_, which were highly correlated (Table 2).

**Table 1.** Marginal posterior distributions of the parameters in the spatio-temporal stochastic aphid population growth model spatial when fitted to 2016 occurrence data. The distributions are summarized by their 2.5%, 50% and 97.5% percentiles.

| Parameter | 2.5% | 50% | 97.5% |
| --- | --- | --- | --- |
| $r_0$ | 0.0147 | 0.0175 | 0.0215 |
| $r_1$ | -1.17E-05 | 1.32E-05 | 2.87E-05 |
| $r_2$ | -1.17E-07 | -8.83E-08 | -4.71E-08 |
| $\sigma_p$ | 0.00554 | 0.00911 | 0.01677 |
| $\mu_0$ | 0.000395 | 0.00786 | 0.01531 |
| $\sigma_0$ | 0.00509 | 0.00857 | 0.01620 |
| $\rho_0^2$ | 0.936 | 1.2185 | 1.2412 |

**Table 2.** Correlation matrix between the parameters *r*_0_, *r*_1_, and *r*_2_ in the spatio-temporal stochastic aphid population growth model spatial when fitted to 2016 occurrence data.

| | $r_0$ | $r_1$ | $r_2$ |
| --- | --- | --- | --- |
| $r_0$ | 1 | -0.96 | 0.93 |
| $r_1$ | -0.96 | 1 | 0.99 |
| $r_2$ | 0.93 | 0.99 | 1 |

The posterior marginal distribution of the parameter that measures the effect of geographic distance on the spatial covariance, *ρ*_0_, is left-skewed towards the upper boundary and significantly larger than zero (Table 1), and the site-specific initial occurrence probabilities are consequently positively correlated among the sites at the spatial scale of 10 km. However, the importance of this positive correlation for the among-site variation in aphid outbreaks has to be evaluated in relation to the estimated among-site variation in the initial occurrence probability as modelled by *σ*_0_, and population growth as modelled by *σ*_*p*_ (Table 1).

The rainfall of the 2016 season is presented in the electronic supplement. Incidents of heavy rain were observed in late May and in June, especially in the Viborg region, with incidents of up to 25 mm per day. However, if the observed rain records at the sites were manually scored according to severity and compared to the latent variables that model the among-site variation in population growth, *ϵ*_*i*_, then there was no significant relationship between the heavy rain score and population growth rate (results not shown, P = 0.89).

The variation in aphid occurrence among sampling plots as a function of the geographical distance among plots on May 30 and June 7 2017 is shown as a semi-variogram (Fig. 5). Generally, the variation among plots is relative low at the two sampling days, although the variation seems to increase irregularly with time when assessed from only two samples in time. There is a slight indication that the variation among plots increases with the distance among plots at the second sampling.

**Fig. 5.**
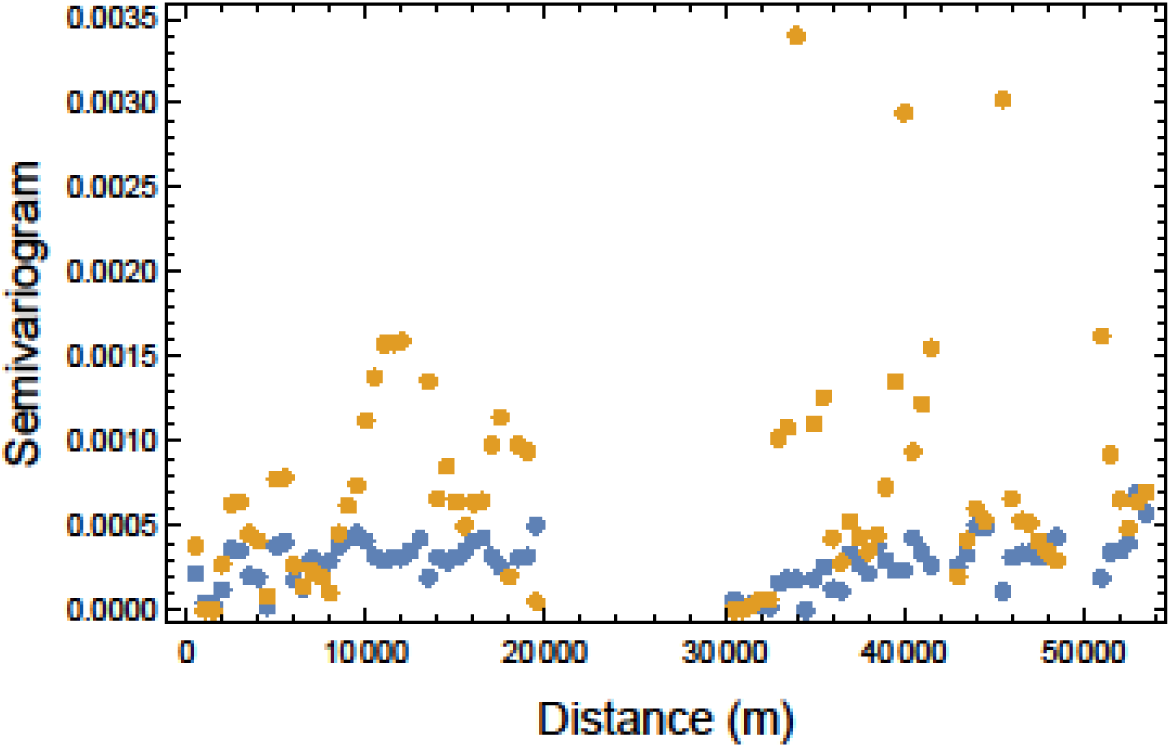
Semivariogram. The variation in aphid occurrence as a function of the geographical distance among sampling plots. Blue points: May 30 2017, yellow points June 7 2017.

## Discussion

The site-specific initial occurrence probabilities were found to be positively correlated among the sites at the spatial scale of 10 km. However, when the spatial variation in the initial occurrence probability was examined in more detail the following year, the spatial variation among plots in the beginning of the aphid outbreaks did not seem to increase much with among-plot distance. This indicates that there was only limited spatial effects, i.e. that the initial epidemic development was more or less in synchrony over distances up to and above 10 km. Since the parameter that measures the effect of geographic distance on the spatial covariance, *ρ*_0_, also depends on the spatial variation in the following aphid epidemic estimated from the 2016 data set, we tend to put more weight on the more detailed investigation in 2017, and conclude that our investigation suggest that there is limited spatial variation in the initial occurrence probability. Consequently, the overall results support the working hypothesis that initial aphid population sizes and outbreaks may be predicted for fields within a 10 km radius of the aphid-monitoring site. For farmers, this may imply that they can rely their decision of whether to spray against aphids on observations made within a distance of up to 10 km.

We did not detect any significant effects of heavy rain events on aphid occurrence probability. This is contradictory to the conclusions of Whitney et al. (2016) and the experimental findings by Mann et al. (1995) that heavy rainfall may cause a loss of aphids up to 30 % of the population. The rainfall incidents measured per day in the present study resemble those studied by Chamuene et al. (2018), who found that such incidents may cause increased aphid mortality and hence reduce population growth in some, but not all cases. One reason why we did not see a reduced population increase may be that most rainfall incidents happened after the ears of the winter wheat appeared, as Jones (1979) only found rainfall before ear appearance to affect aphid numbers.

The simple quadratic function used to model population growth as a function of degree-days performed adequately for modelling the growth in the aphid population from June 1 2016 to July 1 2016. However, the purpose of this simple quadratic model was only to model the deterministic part of the population growth in order to quantify the stochastic variation among sites, and the fitted quadratic model is not suitable for making actual predictions outside the domain of the collected aphid population data. Instead, the deterministic part of the population growth could be the models described by Holst et al. (1997), Ciss et al. (2014), Duffy et al. (2017) or the model element of the aphid modules of decision support systems such as CPO (Hagelskjær & Jørgensen 2003) and GETLAUS (Gosselke et al. 2001).

Most decision support systems will let the user know if it is appropriate to treat against aphids or not, without telling how certain the “decision” is (e.g. GETLAUS (Gosselke et al. 2001) and CPO (Hagelskjær & Jørgensen 2003). If the output came with uncertainties, such as “It can now with 55% certainty pay off to treat against aphids in your field”, using the decision support system may appear more difficult, because the farmer is left with an uncertain foundation for his decision. However, the output might include information on how to reduce the uncertainty. If the farmer had counted aphids on 50 tillers, the output might tell the farmer that the uncertainty would be reduced if he continued and counted on a higher number of tillers, and he could continue until he was able to take certain decisions. This would require some time, but the farmers would get some knowledge on the importance of spending time on providing the decision support system with good initial aphid density estimates, as spending more time will provide a safer foundation for decisions. Estimates of initial densities can be used to predict peak densities of *S. avenae* (Honek et al. 2017), and good estimates of initial densities will, everything else equal, provide better estimates of peak densities.

Nevertheless, the farmer might still not be willing to spend the required time on counting aphid, but counting might be found worth the effort if the result could be used by neighbors, or by all farmers within a certain area. This would optimize the balance between time input spend on counting aphids and economic output in terms of higher yield and probably less expenses on pesticide applications.

## Electronic Supplement

Rainfall at the three study sites in 2016

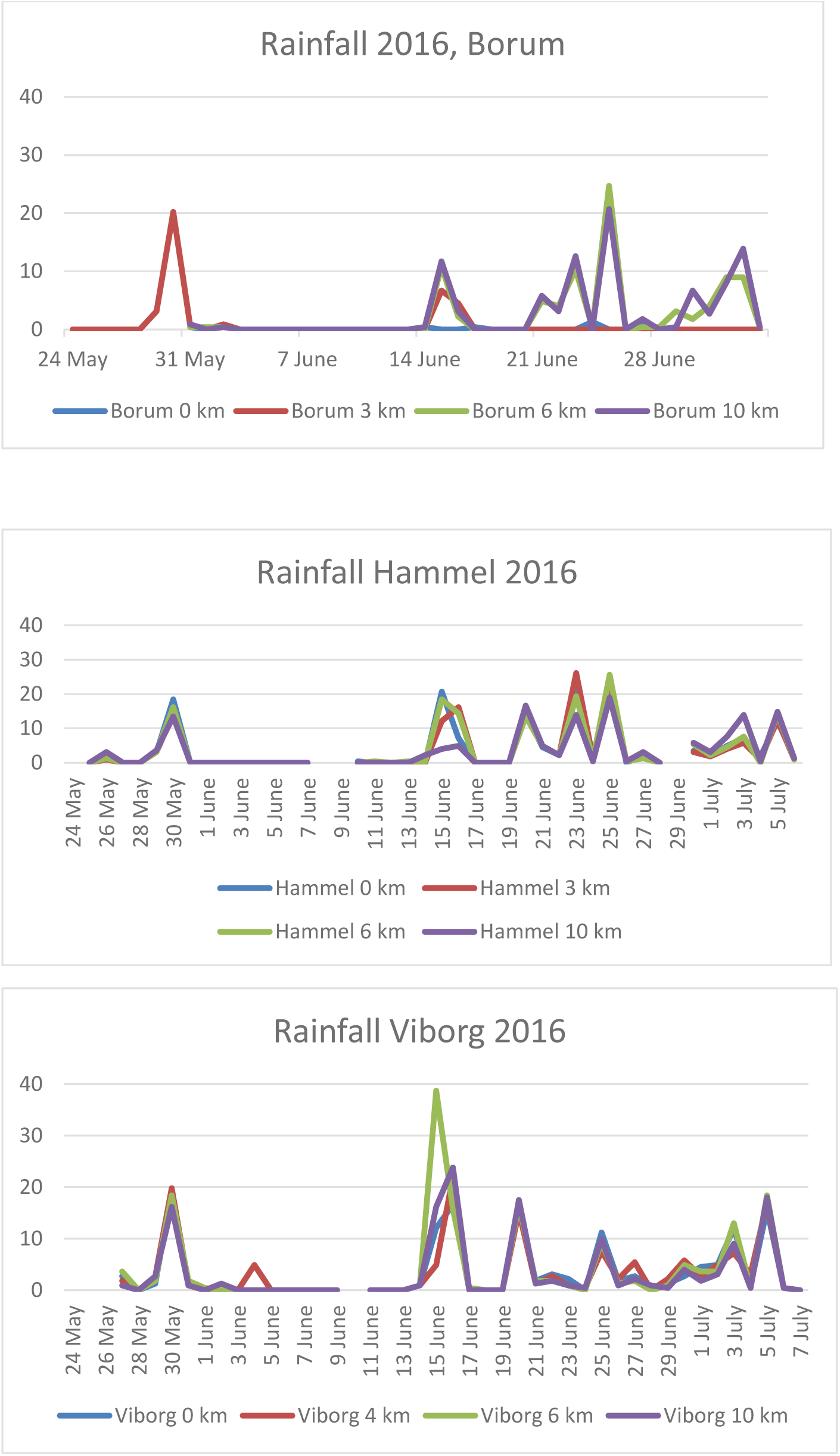

